# Vascular origins of low-frequency oscillations in the cerebrospinal fluid signal in resting-state fMRI: Interpretation using photoplethysmography

**DOI:** 10.1101/2020.10.02.323865

**Authors:** Ahmadreza Attarpour, James Ward, J. Jean Chen

## Abstract

Slow and rhythmic spontaneous oscillations of cerebral blood flow are well known to have diagnostic utility, notably frequencies of 0.008-0.03 Hz (B-waves) and 0.05-0.15Hz (Mayer waves or M waves). However, intracranial measurements of these oscillations have been difficult. Oscillations in the cerebrospinal fluid (CSF), which are influenced by the cardiac pulse wave, represent a possible avenue for non-invasively tracking these oscillations using resting-state functional MRI (rs-fMRI), and have been used to correct for vascular oscillations in rs-fMRI functional connectivity calculations. However, the relationship between low-frequency CSF and vascular oscillations is unclear. In this study, we investigate this relationship using fast simultaneous multi-slice rs-fMRI coupled with fingertip photoplethysmography (PPG). We not only extract B-wave and M-wave range spectral power from the PPG signal, but also derive the pulse-intensity ratio (PIR, a surrogate of slow blood-pressure oscillations), the second-derivative of the PPG (SDPPG, a surrogate of arterial stiffness) and heart-rate variability (HRV). The main findings of this study are: (1) signals in different CSF regions (ROIs) are not equivalent in their vascular contributions or in their associations with vascular and tissue rs-fMRI signals; (2) the PPG signal contains the highest signal contribution from the M-wave range, while PIR contains the highest signal contribution from the B-wave range; (3) in the low-frequency range, PIR is more strongly associated with rs-fMRI signal in the CSF than PPG itself, and than HRV and SDPPG; (4) PPG-related vascular oscillations only contribute to < 20% of the CSF signal in rs-fMRI, insufficient support for the assumption that low-frequency CSF signal fluctuations directly reflect vascular oscillations. These findings caution the use of CSF as a monolithic region for extracting physiological nuisance regressors in rs-fMRI applications. They also pave the way for using rs-fMRI in the CSF as a potential tool for tracking cerebrovascular health through, for instance the strong relationship between PIR and the CSF signal.

## Introduction

Low-frequency hemodynamic oscillations have long been known to exist in the human vasculature (Mayer, 1876; Traube, 1865). Slow and rhythmic spontaneous oscillations of cerebral and peripheral blood flow occur within frequencies of 0.008-0.03 Hz (B-waves) and 0.05-0.15Hz (Mayer waves or M waves). While the generators and pathways of such oscillations are not fully understood, these rhythms have been recognized for their diagnostic significance (Schytz et al., 2010; Spiegelberg et al., 2016). B waves are defined as short repeating elevations in intracranial pressure (Lundberg, 1960; Martinez-Tejada et al., 2019) that can lead to oscillations in arterial blood pressure (Droste and Krauss, 1999), while M waves are interpreted as the magnitude of blood-pressure (BP) oscillations that can translate into blood flow oscillations. An additional complication is that M waves overlap in frequency with vasomotion. M waves are driven by BP oscillations (Julien, 2006; Rieger et al., 2018), and are expected to be systemically synchronous. On the other hand, vasomotion is regionally specific and is defined as the oscillation in vascular tone, which may or may not be accompanied by changes in vascular diameter, but gives rise to flowmotion (Sassaroli et al., 2012), but is not necessarily associated with fluctuations in blood pressure.

The detection of the B wave in gradient-echo echo-planar imaging (EPI) data in the cerebrospinal fluid (CSF) provides a novel non-invasive way to monitor intracranial pressure (C. Strik et al., 2002). CSF flow has also been linked to oscillation in glymphatic flow, through which waste products are removed from the extracellular space by exchange between interstitial fluid and CSF along the perivascular spaces. Pulsation and flow of the CSF are closely linked to the changes in blood flow induced by the heartbeat and by respiration, making pulse and respiration the main drivers of glymphatic flow. However, the role of low-frequency vascular oscillations (~0.1 Hz), specifically vasomotion (which overlaps with the frequency of M waves), has also been reported recently (Marco et al., 2015; van Veluw et al., 2020). CSF oscillations in the 0.001-0.1 Hz range has also been associated with sleep cycles (Fultz et al., 2019). To come full circle, in resting-state functional MRI (rs-fMRI) analysis, the CSF signals (mainly from the ventricles) are routinely taken as a surrogate of cardiac pulsation, and used in nuisance regression when computing functional connectivity, which is primarily based on signals found below 0.1 Hz. The ubiquity of blood-oxygenation level dependent (BOLD) rs-fMRI acquisitions also provide an opportunity to study CSF dynamics and their relationship with other vascular oscillations, potentially allowing the expansion of rs-fMRI for non-invasive intracranial and vascular oscillations.

In the context of rs-fMRI studies, low-frequency vascular oscillations directly overlap with the desired neurogenic signal, and constitute a major source of confound (Chang et al., 2009; Golestani et al., 2017; Tong et al., 2019). The frequency range of B waves overlaps with that of respiratory-volume variations (RVT) (Birn et al., 2006). Moreover, the concern over the interpretation of the 0.1 Hz vascular oscillation stems from observations reported based on optical data (Obrig et al., 2000; Yücel et al., 2016). Given the nuances in the physiological interpretation of different types of low-frequency vascular oscillations and their common frequency range within the rs-fMRI-relevant band, it is of interest to understand how each of the waveforms (B wave, M wave/vasomotion) is associated with the rs-fMRI signal. Many studies have been conducted to find the generators and sources of the low and very low oscillations of MRI data; to that end, time and frequency-domain analysis such as correlation, Fourier and Wavelet transforms have been widely used in the literature (He et al., 2018; Pfurtscheller et al., 2017; Whittaker et al., 2019). Nonetheless, given that these oscillations are difficult to segregate from neuronally-relevant signals by frequency or amplitude, we are unable to fully characterize the origins of these signals. As fMRI signal oscillations in the CSF are naturally not neurogenic, it is widely assumed that they represent a multitude of cardiac-related oscillations which are to be removed from the rs-fMRI data for the purpose of functional connectivity mapping. However, there has yet to be detailed characterization of the vasogenic origins of the CSF signal in rs-fMRI.

The first attempts to extract vascular oscillations from CSF fMRI data were by Strik et al. (Claudia Strik et al., 2002; C. Strik et al., 2002). It was found that changes of arterial blood volume have a major influence on CSF flow, as well as on oscillations of the brain parenchyma (Claudia Strik et al., 2002). In the studies by Strik et al., non-gated resting-state fMRI (rs-fMRI) signal acquired at 1.5 Tesla with rapid sampling (TR of 150 ms) and in a mid-sagittal orientation positioned at the cerebral aqueduct was used to show that while the fast variation of the heart cycle is clearly visible in CSF- and blood flow, whereas slower waves are only detectable in the venous blood flow and heart rate variability. Specifically, B and M wave peaks, while detectable in the CSF and arterial blood, are more pronounced in the venous blood flow (C. Strik et al., 2002). The measurement of the B wave was replicated using EPI MRI data (Friese et al., 2004), taken from a single axial slice positioned perpendicular to the cerebral aqueduct at 1.5 Tesla; again, high sampling rate (repetition time = 110 ms) was used for this purpose. In this latter study, the MRI signal spectrum was separately into 4 pre-defined segments identified in earlier studies (Lang et al., 1999; C. Strik et al., 2002). Thus, it was shown that using fast EPI-MRI acquisitions, slow rhythmic oscillations can be analysed non-invasively using CSF flow captured by rapid fMRI.

Furthermore, from a clinical perspective, there is increasing recognition of the PPG signal as a means to provide cuffless measures of vascular health (Attarpour et al., 2019; Kanders et al., 2013). PPG-derived diastolic pressure (DBP) agrees with that obtained using brachial oscillometry (although PPG-based systolic blood pressure (SBP) does not) (Allan et al., 2018). More recently, low-frequency PPG variance has been suggested to provide a non-invasive estimation of blood pressure, either through simple signal modeling (Sharma et al., 2017) or through machine learning (Attarpour et al., 2019). The PPG intensity ratio (PIR) has been found to be a plausible reference for M-wave-like vascular-diameter oscillations (Ding et al., 2017; Ding and Zhang, 2015). Also, the second deviation of PPG (SDPPG) signals is highly correlated with arterial compliance and stiffness. Thus, not only is there value in identifying the vascular origins in the CSF signal fluctuations, but also in leverage this knowledge generate markers of vascular health from the CSF signal.

In this work, we analyze the dynamics of CSF flow as captured using fast rs-fMRI. Based on the aforementioned previous works, we hypothesize that information about vascular oscillations can be observed in resting-state fMRI data (typically acquired for functional-connectivity mapping), so long as the data sampling rate is sufficiently high. In this work, our primary goal is to investigate the extent to which these various vascular oscillations are found in the rs-fMRI signal of the CSF, and to relate the findings to the same vascular contribution in the rs-fMRI signal from the vasculature (arteries and veins) and brain parenchyma. Moreover, we for the first time also examine the association between the rs-fMRI signal and PPG-derived metrics in not only the CSF, but also in the vasculature and brain tissue. This knowledge will enable the use of the widely available rs-fMRI data for vascular monitoring in addition to its conventional functional-network mapping, as well as inform the efforts in physiological denoising of rs-fMRI signal.

## Methods

### MRI Acquisition

MRI data were collected from 15 healthy adults (mean age 30 ± 6.7 years) on a 3T Siemens TIM Trio scanner and a 32-channel head coil. Specifically, whole-brain resting-state fMRI (rs-fMRI) data were acquired using using simultaneous multi-slice (SMS) acceleration on the gradient-echo echo-planar imaging (EPI) with leak-block slice GRAPPA recon with a 3×3 kernel (Cauley et al., 2014) (TR=380 ms, TE = 30 ms, flip angle= 40°, 15 slices, 3.44×3.44×6.0 mm^3^, 2230 time points, acceleration factor=3, phase encoding shift factor=2, slices ascending). A 3D T1-weighted anatomical scan was acquired using MPRAGE, with resolution 1 × 1 × 1 mm, repetition time (TR) = 2400 ms, inversion time (TI) =1000 ms, echo time (TE) = 2.43 ms, flip angle = 8°, field of view = 256 × 256 mm (sagittal), matrix size = 256 × 256, 192 slices (ascending order), bandwidth = 180 Hz/pixel, and GRAPPA acceleration factor = 2.

During the fMRI scans, Cardiac pulsation was recorded using the Siemens scanner pulse oximeter (sampling rate = 50 Hz), whereas the respiratory signal was recorded using a pressure-sensitive belt connected to the Biopac™ (Biopac Systems Inc. California) at a sampling rate of 200 Hz. The cardiac and respiratory signals were low-pass filtered to 2 Hz and 1 Hz, respectively using a Butterworth infinite-impulse response (IIR) filter with the corresponding bandwidths. Then, each signal is normalized by subtracting the mean and dividing by the standard deviation. Cardiac and respiratory frequencies are estimated as the peak frequencies of the spectra of the filtered (infinite-impulse response) and normalized cardiac and respiratory signals, respectively.

### PPG-derived vascular measures

To help interpret the signals found in rs-fMRI in the CSF, we referenced the rsfMRI signal against features of the PPG signal. These derivatives have different frequency content and physiological interpretations, including:

- B wave: calculated using PPG: B waves were defined as short repeating elevations in intracranial pressure (ICP) (10–20 mmHg) with a frequency of 0.5–2 waves/min (Lundberg, 1960), which translates to a fairly broad range of 0.008 to 0.03 Hz. One way to subdivide B waves is in terms of symmetrical, asymmetrical and plateau waves, each potentially associated with distinct etiology (Martinez-Tejada et al., 2019). Although clearly visible B waves are typically a feature of disease, the frequency of B waves is immediately overlapping with the range of interest for rs-fMRI, prompting us to study B-wave contribution in the CSF signal.
- Low-frequency PPG, a surrogate for the M wave: The M wave is known to represent the 0.1 Hz fundamental oscillation of mean arterial pressure. It is one of the vascular signatures that can be derived from peripheral PPG through spectral analysis (Kanders et al., 2013)
- Heart rate variability (HRV), a surrogate of autonomic nervous regulation: HRV is defined as the time or the number of samples between two consecutive systolic peaks in the ECG or PPG signals. It is dependent only on PPG peak-to-peak distances, not on PPG fluctuation amplitude. It is commonly used in the studies that have been conducted to find non-neuronal sources of oscillations in rs-fMRI data. The results of the studies showed significant correlations between HRV and fMRI data (C. Strik et al., 2002). Kiselev et al. found that LF oscillations in PPG that lag LF HRV is likely to have a neurogenic nature, whereas HRV changes driven by LF PPG changes are likely to have a hemodynamic nature (due to cardiac output) (Kiselev et al., 2020). Quantitative estimation of the degree of synchronization between the LF oscillations in HRV and PPG has potential clinical importance, for example, for assessing cardiovascular risk and controlling drug therapy (Kiselev et al., 2020).
- Derivatives of PPG: Within this category, the second derivative of the PPG signal (SDPPG), also called the acceleration PPG, is much more commonly used than the first derivative.

- The SDPPG was first proposed by Takazawa et al. (Pilt et al., 2013; Takazawa et al., 1998) as a surrogate of arterial compliance and stiffness. This is a measure that incorporates both PPG peak-to-peak distance and amplitude. SDPPG was computed on PPG data that was filtered with a 4th order Butterworth low-pass filter with a cut-off frequency of 4 Hz.
- PPG intensity ratio (PIR), a surrogate of slow variations in blood pressure: The PIR is a reference for M-wave-like vascular-diameter oscillations (Ding et al., 2017; Ding and Zhang, 2015). As PIR is the ratio of PPG peak intensity to PPG valley intensity of one cardiac cycle, it is highly dependent on PPG fluctuation amplitude and not on beat-to-beat distance.

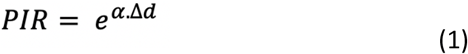

where *α* is considered to be a constant related to the optical absorption coefficients in the light path, and Δ*d* reflects the arterial diameter change during one cardiac cycle from systole to diastole. It has been demonstrated that the variations of PIR were associated with those of SBP during deep breathing and the Valsalva manoeuvre, with PIR correlated negatively with SBP during deep breathing but positively during the Valsalva manoeuvre. This is justified by the reduction in sympathetic nervous activity during deep breathing that leads to the relaxation of vascular smooth muscle, and further the increased arterial diameter and elevated PIR accordingly (Sharma et al., 2017). PIR was also computed on PPG data filtered to retain up to 0.4 Hz, as in the case of the SDPPG.

### PPG Data Processing

Firstly, the PPG data were aligned with the fMRI time-series so that their starting points were the same. After implementing a 4th-order Butterworth low-pass filter with the 4 Hz cut-off frequency on the aligned PPG data, SDPPG, HRV, and PIR were obtained. PIR and HRV signals were firstly interpolated using the Cubic spline data interpolation method to have an equal length with the PPG data. Subsequently, for further analysis, the PPG data in addition to SDPPG, HRV, and PIR had to be resampled to have an equal length with fMRI time series. Before resampling, a first-order Butterworth low-pass filter with 1 Hz cut-off frequency was used as an anti-aliasing filter.

In all statistical analyses in this study, the data were firstly filtered to 0.005 to 1.2 Hz (except PPG data in the Frequency Components analysis that were filtered to 0.005 to 4 Hz) using the 4th-order Butterworth filters and then were centered around mean with a unit standard deviation.

Power spectral density (PSD) of all the fMRI ROIs and PPG-driven data were calculated using Welch’s overlapped segment averaging estimator. For this analysis, a 250 sec Hamming window with a 90% overlap was used to cover two cycles of B waves. As a result, the lowest frequency that can be considered is 0.004 Hz. The percentage of B wave and M wave/Vasomotion’s frequency bands of each signal was determined by dividing the integral of each band of frequency to the integral of the whole PSD. For comparison of the results with other studies, the values of mean and standard error among all signals and subjects were subsequently computed.

### Image Data Preprocessing

For all subjects, FreeSurfer reconstruction was performed on the T1 anatomical data using FreeSurfer 6.0 (publicly available: https://surfer.nmr.mgh.harvard.edu). The reconstruction provided tissue segmentation of grey matter, white matter structures as well as ventricles, which are used later for delineating the regions of interest.

The rs-fMRI processing pipeline includes: motion correction, spatial smoothing (Gaussian kernel with 5mm FWHM), high-pass filtering (> 0.001 Hz), and registration of data into a 4×4×4 mm^3^ MNI atlas (45 × 54 × 45 voxels). All steps were performed using FMRIB software library (FSL, publicly available at www.fmrib.ox.ac.uk/fsl). As we aim to characterize the potential effects of vascular pulsation in rs-fMRI data, we did not regress out nuisance variables in the preprocessing stage.

### Regions of Interest

As the primary aim of this work is to investigate the rs-fMRI signal fluctuations in the CSF, we defined several disjoint CSF-related regions of interest (ROIs):

- Lateral ventricles and third ventricle: Both of these structures are taken from the FreeSurfer tissue segmentation, and subsequently resampled to the spatial resolution of the rs-fMRI images using FSL flirt. The bilateral lateral ventricles (LV) are both included in the LV ROI.
- Cerebral aqueduct: This is the smallest of all visible CSF structures, and was delineated manually in the rs-fMRI images using the T1 anatomical image as reference.

Other ROIs were also defined for reference, as some of them (such as blood vessels) have been linked with CSF pulsations, and others (brain tissue) have been reported to also contain residual CSF pulsatility:

- Arteries: First, the rs-fMRI signal variance map was generated from the pre-processed data. The blood vessels have consistently high variance (Churchill and Strother, 2013), thus the variance maps were used for delineation of blood vessels. For each subject, the mask for arteries were delineated manually based on known vascular anatomy, made to contain the Circle of Willis as well as branches of the internal carotid arteries and middle-cerebral arteries.
- Veins: As in the case of the arteries, veins were also delineated manually. The mask for veins includes the superior sagittal sinus and the transverse sinus.
- Grey matter (GM) and white matter (WM): The GM and WM masks were both delineated on a per-subject basis using the tissue segmentation results produced by the FreeSurfer recon process. These masks were also subsequently resampled to the spatial resolution of the rs-fMRI images using FSL flirt.

With the exception of the aqueduct, all ROIs are eroded by 1 voxel to minimize partial-volume effects.

### Statistical analysis

#### Correlation Analysis

To find out the similarity and the relationship between different fMRI ROIs, the Pearson correlation coefficient and its associated p-value were calculated. If a determined p-value was smaller than the significance level (0.05), then the corresponding correlation was considered significant. Finally, the averages of the absolute correlation coefficients among all subjects were computed.

#### Cross Correlation Analysis

In this step, the fMRI time-series and PPG-driven data were firstly filtered to 0.008-0.03 Hz (B wave) using the 4th-order Butterworth filters. As the fMRI signals are non-stationary, sliding-window cross-correlation between each fMRI time-series and PPG, HRV, SDPPG, and PIR was calculated. Each signal was divided into 250-sec sections with a 50% overlap to cover two cycles of B waves and the final cross-correlation result was obtained by averaging along time. Subsequently, the afore-mentioned procedure was repeated for analyzing the M wave (0.05-0.15 Hz). In this study, positive time lags represent that fMRI time-series lead PPG data.

Moreover, for each run, the maximum positive correlation coefficient at positive time lags (when data were filtered to B wave frequency range) and at negative time lags (when data were filtered to M wave/vasomotion frequency range) was computed. As each subject may have a different time lag, the mean of the maximum positive correlation coefficient and the lags corresponding to those maximum positive correlations among all subjects were calculated.

#### Cross Spectrogram

Cross spectrograms between each fMRI time series and the corresponding PPG, HRV, SDPPG, and PIR were calculated by using the Short Time Fourier Transform (STFT). The cross spectrogram between two signals highlights the frequencies that they have in common. For that purpose, a 125 sec Hamming window with a 90% inter-step overlap was used that results in having frequency components higher than 0.008 Hz. In our study, one constraint had to be considered was considering the frequency of higher than 0.008 Hz, and that lead us to choosing the 125-sec window for this analysis. In the STFT analysis, a trade-off must be considered between time and frequency domains, and the choice of 125 s window length reflects that trade-off.

The maximum-energy time-frequency ridge was then determined by averaging three obtained ridges weighted by the amplitude of the cross-spectrogram results. For example, the final ridge between the LV time series and PPG data represents the most common frequencies over the time period that these two signals have in common. By averaging the frequencies of the final ridge, the main common frequency between the two signals was computed. All the above-mentioned cross spectrogram analyses were repeated three times when only B wave frequency range (0.008-0.03 Hz), M wave frequency range (0.05-0.15 Hz), and frequency range of 0.008-1 Hz was selected as the frequency range of interest.

## Results

Of the 18 subjects, 3 had to be excluded due to failure to critically sample the fundamental cardiac frequency (as the 2 subjects had heart rates higher than 1.3 Hz). Furthermore, data from 4 of the subjects did not allow for inclusion of the cerebral aqueduct.

In Figure 2 we show the regional-average rs-fMRI time series plotted for CSF regions as well as vascular and tissue ROIs for comparison. It can be observed that the rs-fMRI signals in the lateral ventricles and third ventricle are synchronized with those in the arteries and veins, and to a lesser extent, those in the grey matter and white matter. Moreover, the signals in the ventricles appear to be better reflected in the PIR than in the SDPPG, the HRV and in the PPG itself.

**Figure 1.**
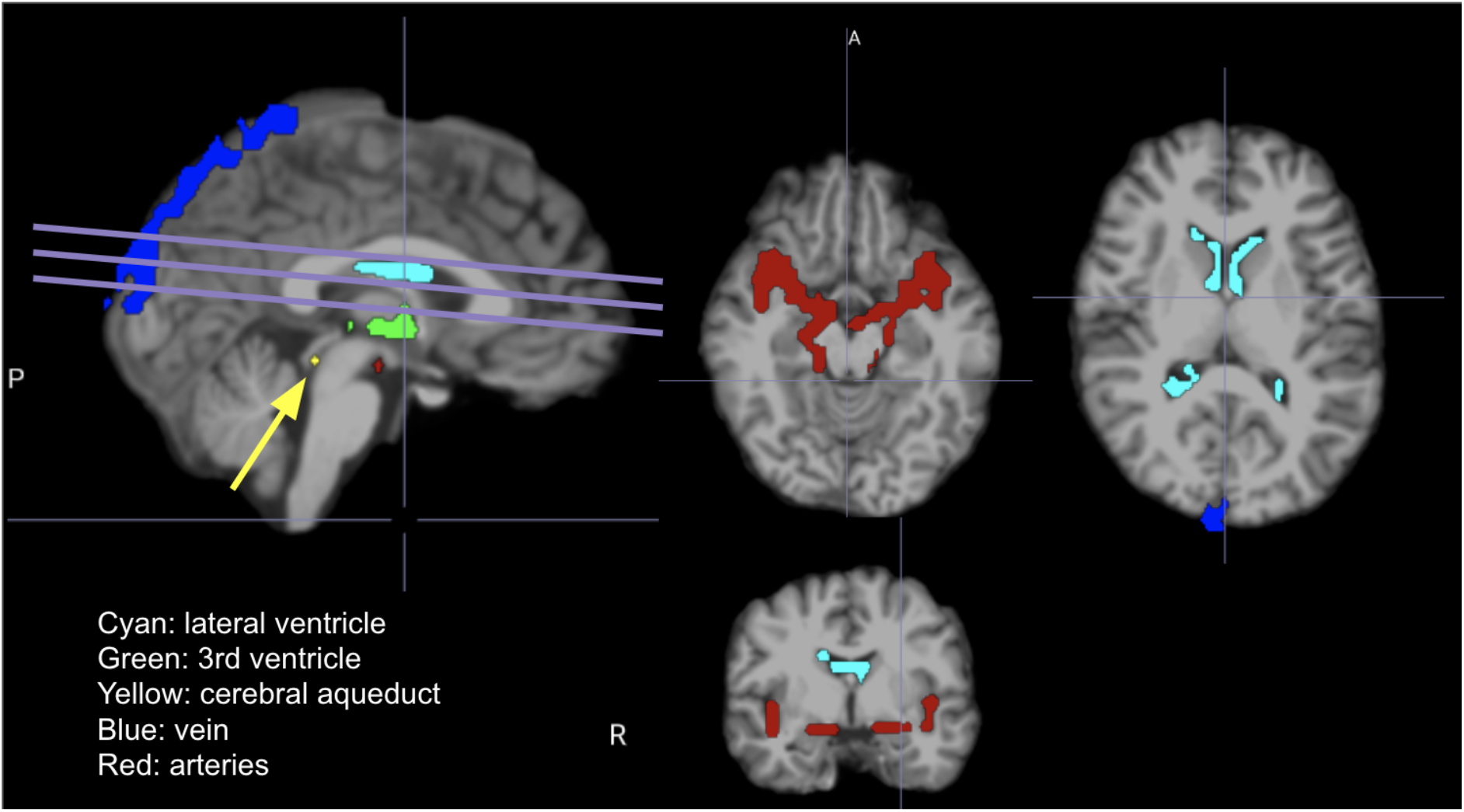
Regions of interest, specifically CSF and vascular ROIs, for a representative subject, overlaid on the T1 anatomical. The lines indicate the fMRI slice orientation, and the arrow indicates the aqueduct. A = anterior, P = posterior, R = right.

**Figure 2.**
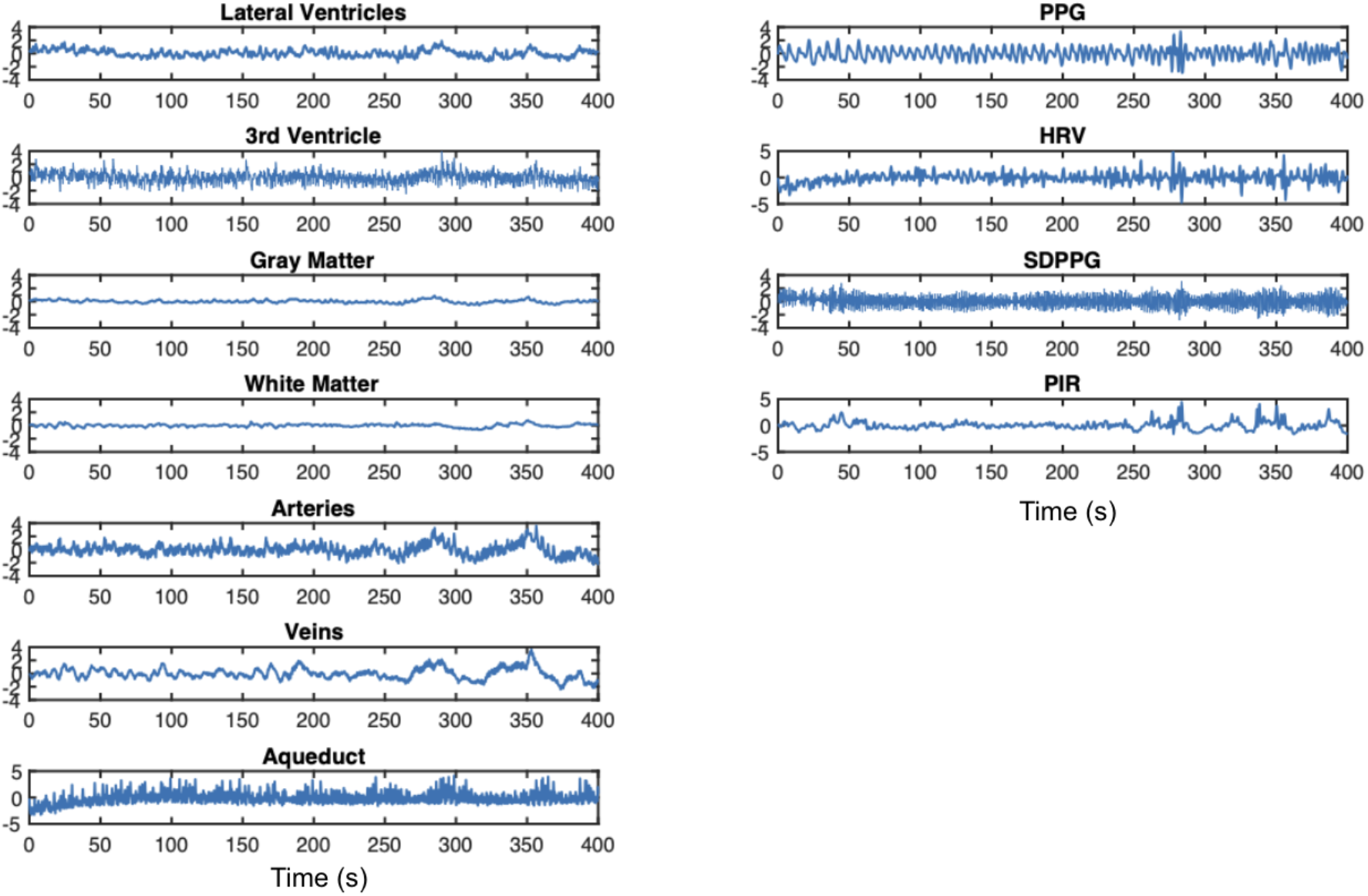
Time series the fMRI in CSF-related ROIs, contrasting signals from other ROIs as well as the PPG-associated signals, from a representative subject. CSF-related ROIs include the lateral ventricles (LV), the third ventricle (3rd V) and the cerebral aqueduct. All signals have been resampled (to the maximum frequency of the rsfMRI data). Column 1: It can be observed that the rs-fMRI signals in the LV and 3rd ventricle are synchronized with those in the arteries and veins, and to a lesser extent, those in the GM and WM. Column 2: It can also be observed that PIR best follows signals in these ROIs.

The time-series patterns are mirrored in the spectral plots in Figure 3. It is evident that the third ventricle and the cerebral aqueduct contain more signal at the cardiac frequency than the lateral ventricles. The aqueduct also contains a peak at the respiratory frequency (0.25 Hz), which is not evident in the other CSF ROIs. Moreover, in terms of PPG-derived features, the spectral characteristics of PIR are more similar to that of the LV and the blood vessels, echoing observations in the time domain (Figure 2). On the other end of the spectrum, SDPPG, like the aqueduct, appears to be mainly driven by high frequencies (heart rate), which is also strongly manifested in the aqueduct fMRI signal.

**Figure 3.**
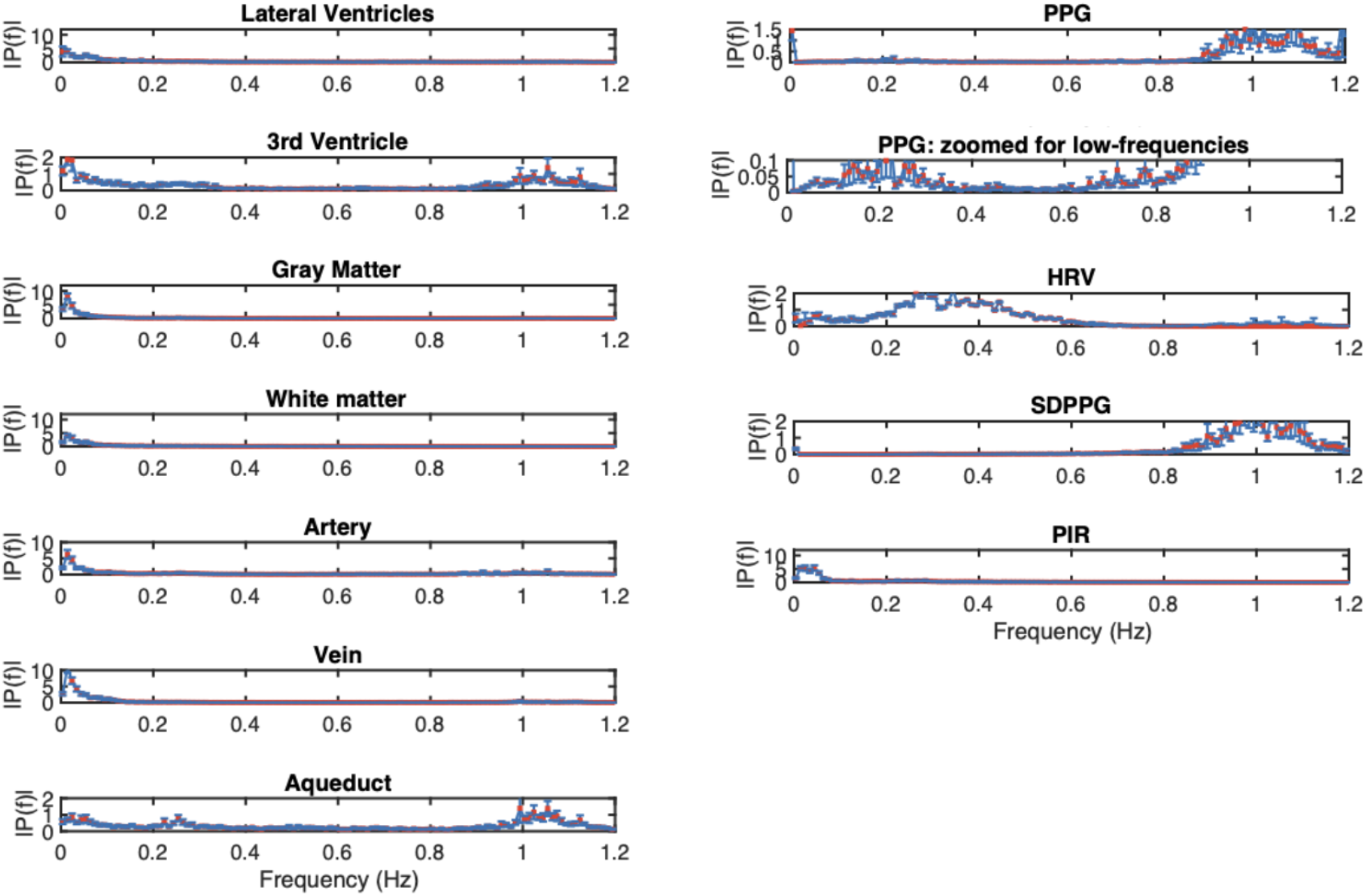
Frequency spectrum of the fMRTime series in CSF-related ROIs, contrasting spectra from other ROIs as well as the PPG-associated spectra. Spectra are averaged across subjects, and error bars represent standard error. CSF-related ROIs include the lateral ventricles (LV), the third ventricle (3rd V) and the cerebral aqueduct. All signals have been resampled (to the maximum frequency of the rsfMRI data). Notice that for the PPG spectrum, the cardiac peak is substantially higher than the low-frequency peak.

These relationships are further clarified in the correlation matrices in Figure 4. In the low-frequency range ([0.008 - 0.15 Hz]), the signals in the lateral ventricles (LV) are significantly correlated with those in the 3rd ventricle (3rd V), large arteries, the grey matter (GM) and the white matter (WM). However, the highest correlations (that are also significant) are not found in the CSF regions. Rather, they are found between WM and GM, between GM and large arteries, between WM and large veins, and between GM and large veins. These patterns are reproduced in the broadband range, namely [0.005 - 1.2 Hz].

**Figure 4.**
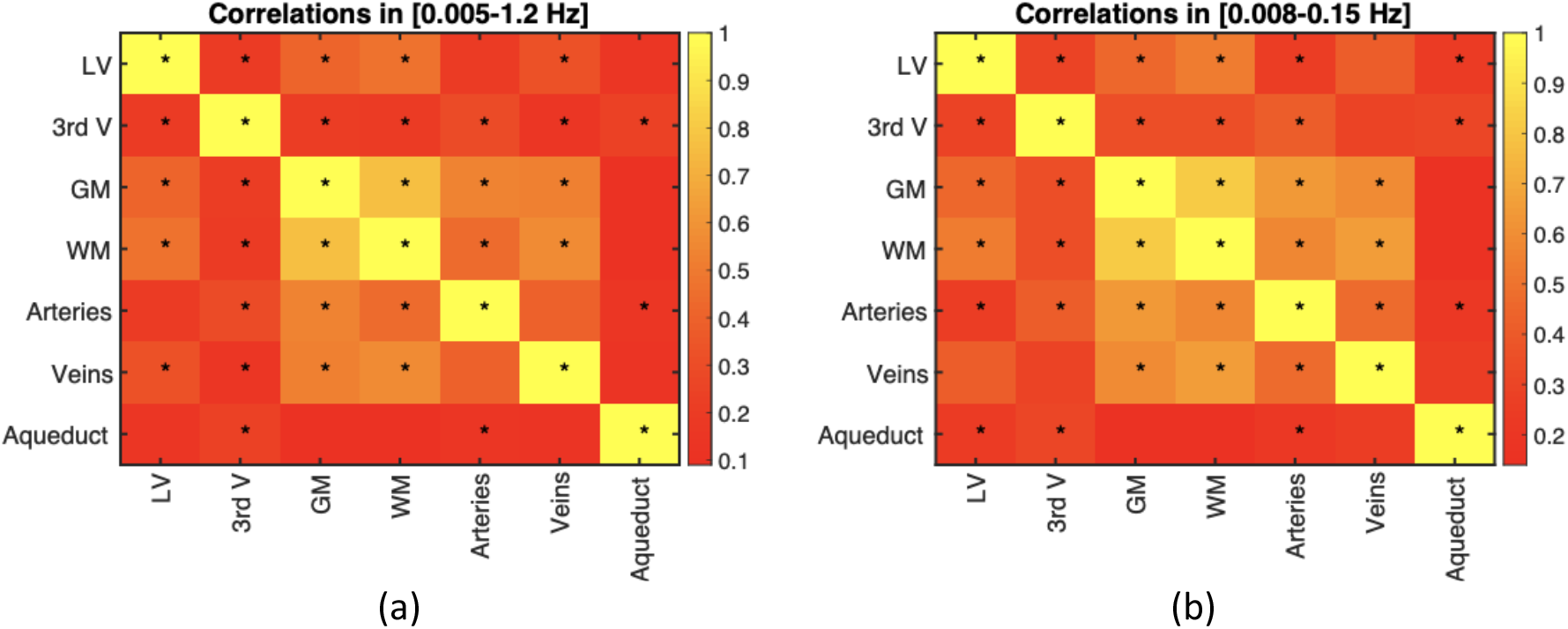
Signal correlation amongst different ROIs. The correlation coefficients are the average across all subjects. Statistical significance of the correlations are indicated by asterisks (*: p < 0.05). The signals in the lateral ventricles (LV) are significantly correlated with those in the 3rd ventricle (3rd V), the grey matter (GM), the white matter (WM) and large arteries. However, the highest correlations (that are also significant) are not found in the CSF regions. Instead, they are found between WM and GM, between GM and large arteries, between WM and large veins, and between GM and large veins. These patterns are reproduced across two frequency ranges, namely [0.005 - 1.2 Hz] (a) and [0.008 - 0.15 Hz] (b).

As can be seen in Figure 5a, when averaged across all subjects, the B-wave frequency range contributes less than 20% of the fMRI signal from CSF regions (lateral ventricles, third ventricle and aqueduct). In comparison, this frequency contributes more to the grey matter ROI, followed by large-vein ROIs, WM and arterial regions. Amongst the PPG-derived measures (Figure 5b), the B-wave frequency range accounts for the most power in the PIR signal, followed by the raw PPG signal, HRV and SDPPG.

**Figure 5.**
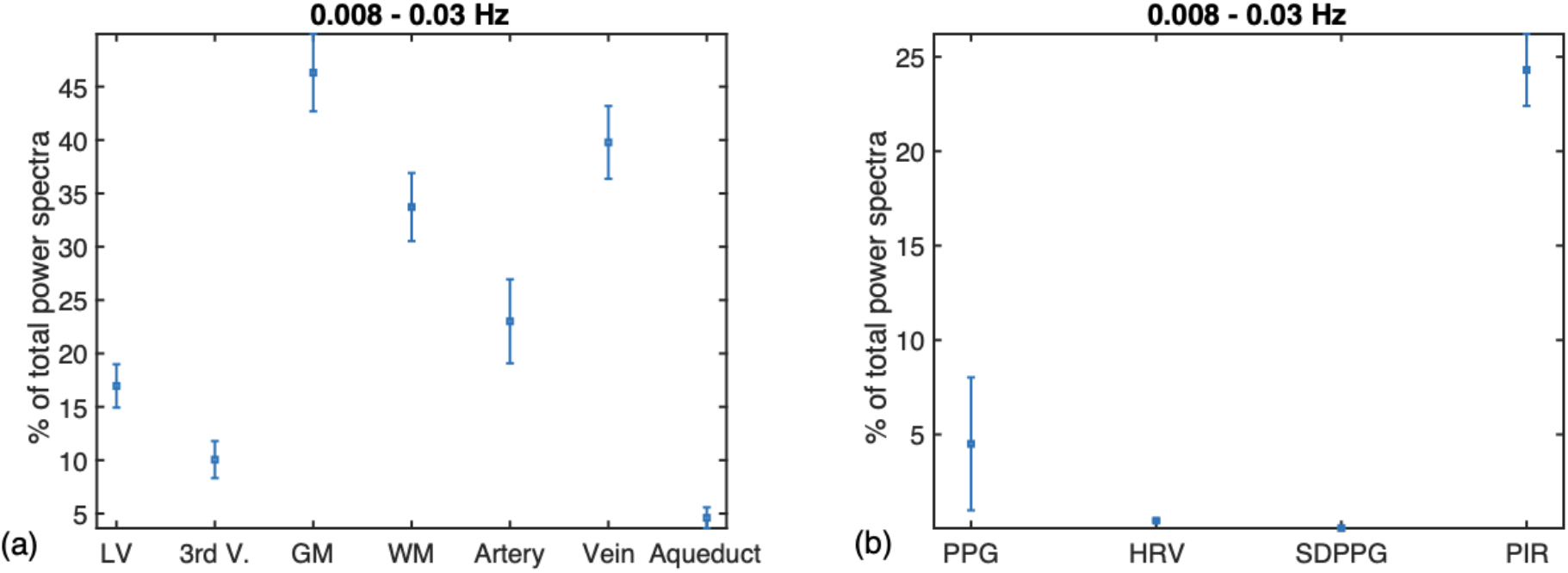
The percent signal power found in the B-wave frequency band. These are presented as fractions of the total spectral power up to 1.2 Hz. (a) On average (across all subjects), the B-wave frequency range contributes < 20% of the fMRI signal from CSF regions (LV, 3rd V and aqueduct). This frequency contributes the most to the rs-fMRI signal in the grey matter (GM), followed by large veins, white matter (WM) and arteries. (b) Amongst the PPG-derived measures, the B-wave frequency range accounts for the most power in the PIR signal, followed by the raw PPG signal, HRV and SDPPG. LV: lateral ventricles; 3rd V: third ventricle; GM: grey matter; WM: white matter. Error bars represent the standard deviation across subjects.

On the other hand, the M-wave frequency range accounts for the highest percent rs-fMRI signal power in the lateral ventricles, followed by the white matter (WM) and large veins (Figure 6a). Amongst the PPG-derived measures, the M-wave frequency range accounts for the most power in the PPG signal, followed by the PIR signal, HRV and SDPPG (Figure 6b).

**Figure 6.**
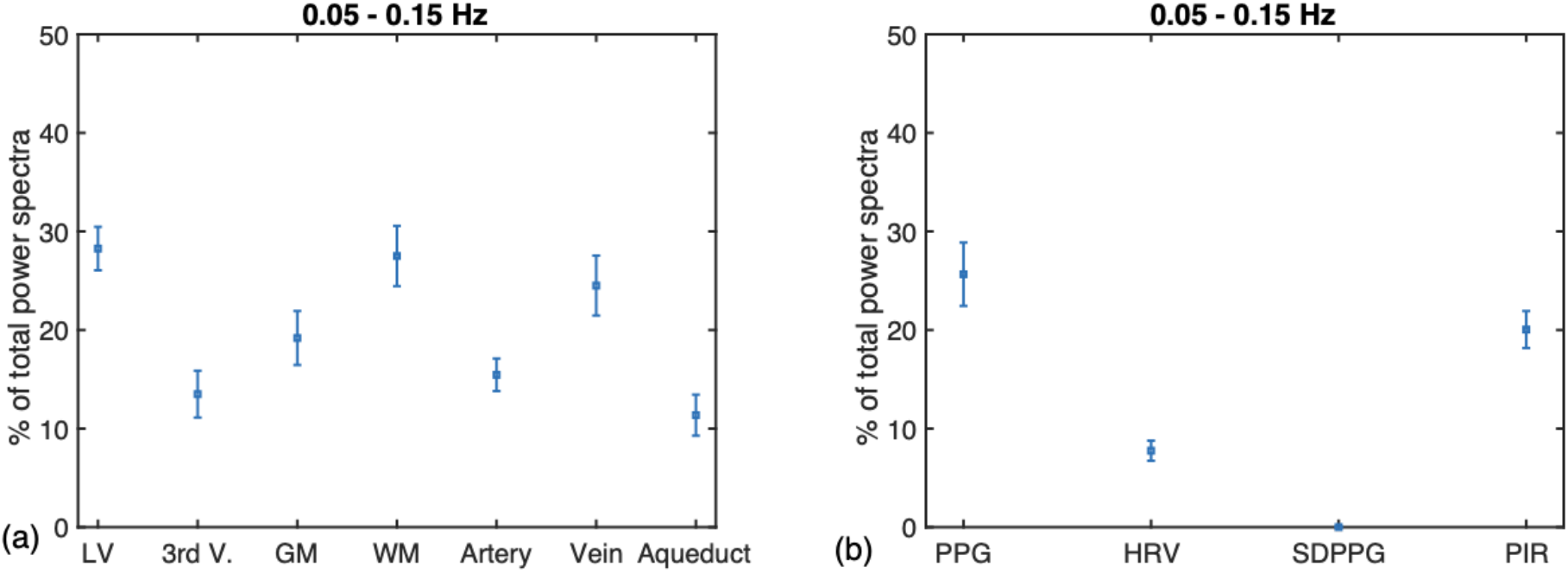
The percent signal power found in the M-wave frequency band. These are presented as fractions of the total spectral power up to 1.2 Hz. (a) On average (across all subjects), the M-wave frequency range contributes most to the rs-fMRI signal in the lateral ventricles (LV), followed by the white matter (WM) and large veins. (b) Amongst the PPG-derived measures, the M-wave frequency range accounts for the most power in the PPG signal, followed by the PIR signal, HRV and SDPPG. LV: lateral ventricles; 3rd V: third ventricle; GM: grey matter; WM: white matter. Error bars represent the standard deviation across subjects.

In the B-wave frequency range (Figure 7), the cross-correlation between the PPG signal and various fMRI ROIs peak at approximately 20 and − 20 s. The same is found for the most part between HRV and fMRI signals (across all ROIs except for the cerebral aqueduct). The strongest correlations between PPG and rs-fMRI signals are found in the lateral ventricles. For SDPPG, the relationship with fMRI signals is more complex; with the LV ROI, the GM and WM as well as arteries and veins, the peak lags are mostly positive, indicating PPG lagging the fMRI signal, but this is not the case in the remaining CSF ROIs. With PIR, the strongest and most reproducible correlation peaks are found in the GM and WM ROIs, with lags close to zero.

**Figure 7.**
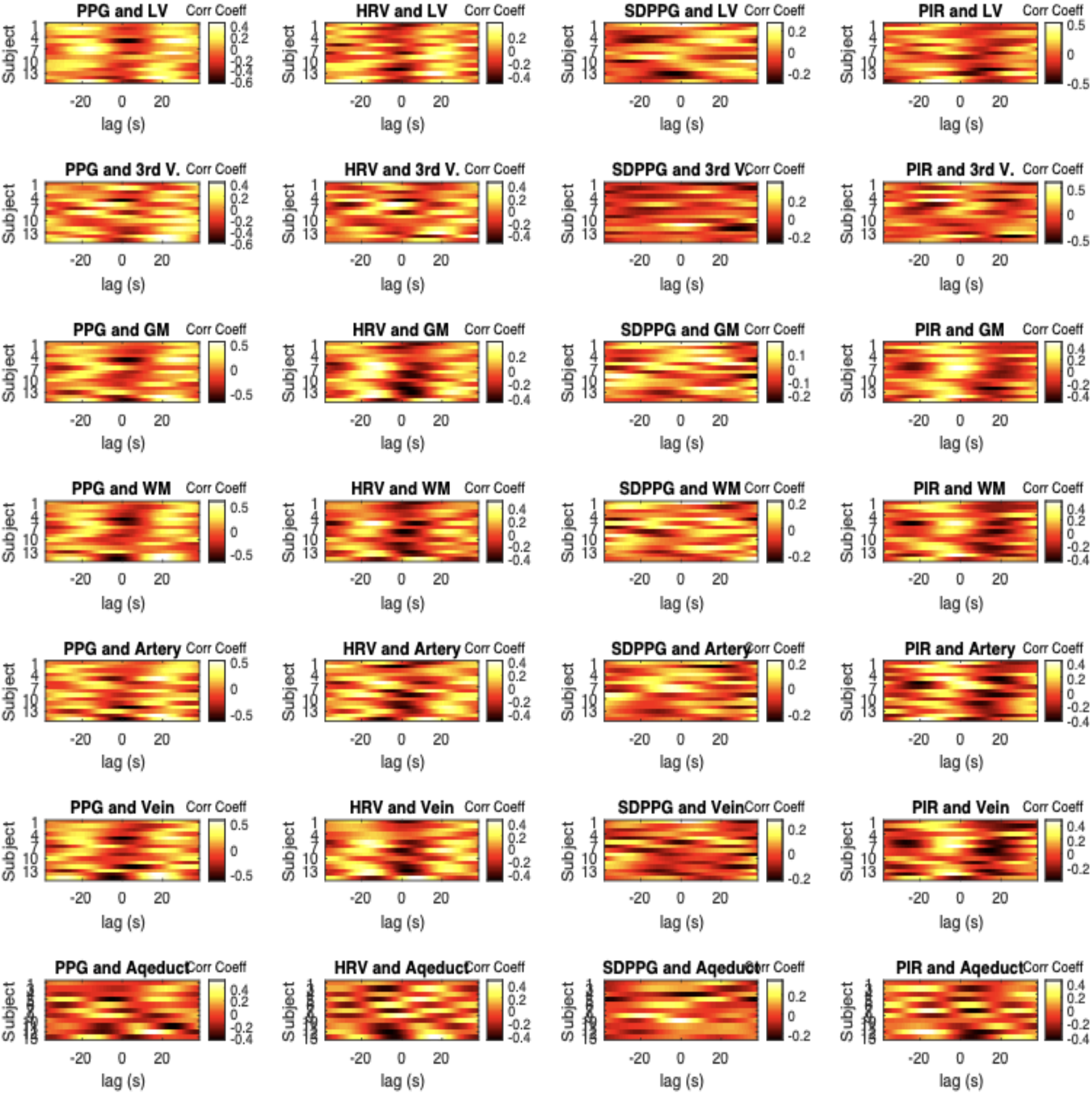
Cross-correlation carpet plot for the B-wave frequency band. In each plot, each row represents the cross-correlation pattern from one subject. Positive lags indicate fMRI leading PPG. Each The fMRI and PPG-derived signals were all bandlimited to 0.008-0.03 Hz. Column 1: In this range, the cross-correlation between the PPG signal and various fMRI ROIs peak at approximately 20 and - 20 s, suggesting a cyclic cross correlation between the two. Column 2: The same is found for the most part between HRV and fMRI signals (across all ROIs except for the cerebral aqueduct). Column 3: For SDPPG, the relationship with fMRI signals is more complex. With the LV ROI, the peak lags are mostly positive, indicating PPG lagging the fMRI signal. Similar patterns are found with the GM and WM ROIs as well as arteries and veins, but not in the remaining ROIs. Column 4: With PIR, the strongest and most reproducible correlation peaks are found in the GM and WM ROIs, with lags close to zero. The remaining ROIs do not share these patterns. LV: lateral ventricles; 3rd V: third ventricle; GM: grey matter; WM: white matter.

Table 1 summarizes the results shown in Figure. 7. As B waves originate as ICP oscillations and propagate into arterial blood pressure (Droste and Krauss, 1999), positive lags are more plausible. In the B-wave range, the mean lag associated with the peak cross-correlations are generally between 14 and 22 s, being the longest for PPG and shortest for PIR. At 0.28-0.34, the cross-correlation of CSF signals with the PPG signal in the B-wave band are comparable to those of the vascular and tissue signals, as are the mean peak correlation coefficients at these lags. However, PIR accounts for the highest cross-correlations with rs-fMRI signals, on average being 0.39 in the lateral ventricles, 0.41 in the third ventricle, and as high as 0.46-0.47 in the grey matter and veins, but the peak cross correlations with PIR are found at negative lags (PIR leading B waves). Note that in general, arteries and the white matter share similar lags, while veins and the grey matter share similar lags with respect to PPG-based signals. Amongst all ROIs, CSF regions also have generally the lowest cross-correlation coefficients with PPG-signals, although the difference is not statistically significant.

**Table 1.** Group-average peak cross-correlation and associated lag for the frequency range 0.008-0.03 hz. All values are listed as mean and standard deviation.

|  | PPG |  | HRV |  | SDPPG |  | PIR |  |
| --- | --- | --- | --- | --- | --- | --- | --- | --- |
|  | Mean peak corr | Mean peak lag (s) | Mean peak corr | Mean peak lag (s) | Mean peak corr | Mean peak lag (s) | Mean peak corr | Mean peak lag (s) |
| **Lateral ventricles** | 0.35±0.03 | 22±1.18 | 0.34±0.02 | 22±1.40 | 0.24±0.02 | 19±1.84 | 0.33±0.02 | 21±1.74 |
| **3rd ventricle** | 0.40±0.03 | 22±1.94 | 0.37±0.02 | 21±2.24 | 0.27±0.03 | 16±1.68 | 0.41±0.02 | 17±2.63 |
| **Grey matter** | 0.41±0.03 | 22±1.54 | 0.31±0.03 | 24±1.31 | 0.19±0.02 | 18±1.98 | 0.41±0.03 | 13±2.02 |
| **White matter** | 0.40±0.03 | 21±1.66 | 0.33±0.02 | 24±1.83 | 0.19±0.02 | 20±1.65 | 0.38±0.04 | 16±1.84 |
| **Arteries** | 0.39±0.03 | 20±1.31 | 0.35±0.02 | 23±1.58 | 0.20±0.01 | 18±1.76 | 0.38±0.03 | 18±1.93 |
| **Veins** | 0.41±0.03 | 22±1.57 | 0.34±0.03 | 24±1.73 | 0.18±0.02 | 19±2.07 | 0.41±0.03 | 16±1.83 |
| **Aqueduct** | 0.35±0.04 | 21±1.67 | 0.41±0.04 | 20±2.09 | 0.27±0.03 | 17±2.11 | 0.35±0.03 | 25±1.53 |

In the M-wave frequency range (Figure 8), there is no dominant lag for the peak cross-correlation between the PPG signal and various fMRI ROIs. For HRV and fMRI, the dominant lag is near −3 s (HRV leading fMRI), observable in all ROIs except for LV. For SDPPG, like in the case of the B-wave frequency, the relationship with fMRI signals is more complex with no visible dominant lag. With PIR, the 3rd ventricle demonstrates a dominant lag at close to 1 sec, which can also be observed in the arterial ROI but not in the other ROIs. Note that signals in LV and 3rd V are more strongly correlated with PIR than with PPG itself.

**Figure 8.**
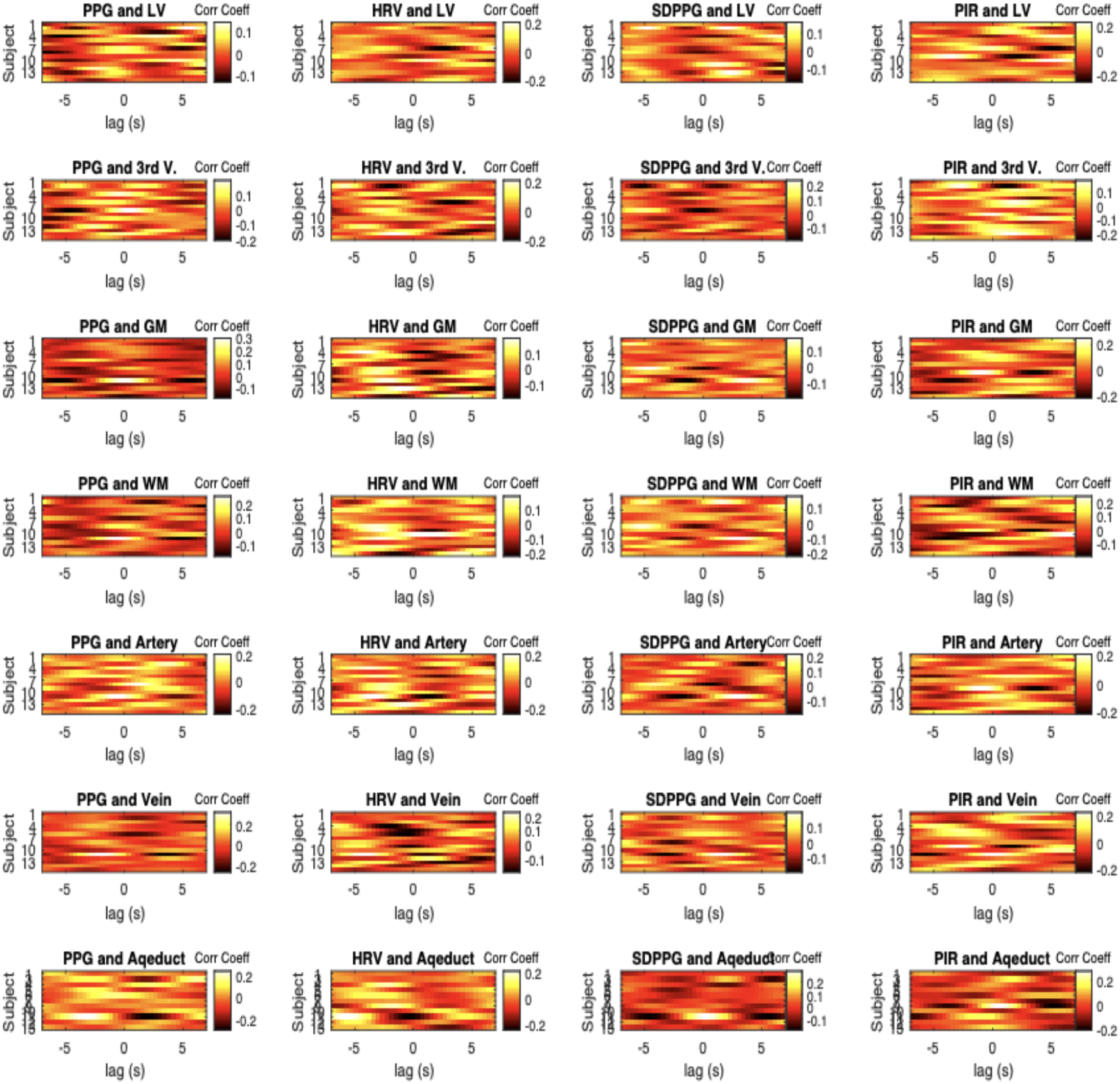
Cross-correlation carpet plot for the M-wave frequency band. In each plot, each row represents the cross-correlation pattern from one subject. Positive lags indicate fMRI leading PPG. Each The fMRI and PPG-derived signals were all bandlimited to 0.05-0.15 Hz. The cross-correlation window has accordingly been reduced from ±20 s to ±7 s. Column 1: In this range, there is no dominant lag for the peak cross-correlation between the PPG signal and various fMRI ROIs. Column 2: For HRV and fMRI, the dominant lag is near −3 s (HRV leading fMRI), observable in all ROIs except for LV. Column 3: For SDPPG, like in the case of the B-wave frequency, the relationship with fMRI signals is more complex with no visible dominant lag. Column 4: With PIR, the 3rd ventricle demonstrates a dominant lag at close to 1 s, which can also be observed in the arterial ROI but not in the other ROIs. LV: lateral ventricles; 3rd V: third ventricle; GM: grey matter; WM: white matter.

Table 2 summarizes the results shown in Figure 8. As before, we focus on negative lags, which indicate the rs-fMRI signal lagging the PPG signal. In the M-wave range, the mean lag associated with the peak cross-correlations are generally between 4 and 5 s, with no noticeable difference among the various PPG-based metrics. At 0.35-0.4, the mean peak cross-correlation of CSF signals with PPG are comparable to those of the vascular and tissue signals, as are the mean peak correlation coefficients at these lags. SDPPG accounts for the lowest cross-correlation values, with no distinction between CSF regions and non-CSF regions. Once again, arteries and the white matter share similar lags, while veins and the grey matter share similar lags with respect to PPG-based signals.

**Table 2.** Group-average peak cross-correlation and associated lag for the frequency range 0.05-0.15 hz. All values are listed as mean and standard deviation.

|  | PPG |  | HRV |  | SDPPG |  | PIR |  |
| --- | --- | --- | --- | --- | --- | --- | --- | --- |
|  | Mean peak corr | Mean peak lag (s) | Mean peak corr | Mean peak lag (s) | Mean peak corr | Mean peak lag (s) | Mean peak corr | Mean peak lag (s) |
| **Lateral ventricles** | 0.40±0.01 | -4.73±0.13 | 0.42±0.01 | -4.91±0.15 | 0.32±0.01 | -4.85±0.15 | 0.41±0.02 | -4.69±0.14 |
| **3rd ventricle** | 0.40±0.01 | -4.57±0.16 | 0.40±0.01 | -4.82±0.19 | 0.35±0.01 | -4.80±0.14 | 0.39±0.01 | -4.74±0.15 |
| **Grey matter** | 0.40±0.01 | -4.75±0.21 | 0.42±0.02 | -4.48±0.21 | 0.31±0.01 | -4.69±0.18 | 0.42±0.01 | -4.51±0.24 |
| **White matter** | 0.39±0.01 | -4.89±0.22 | 0.41±0.02 | -4.70±0.19 | 0.31±0.01 | -4.67±0.17 | 0.41±0.01 | -4.50±0.21 |
| **Arteries** | 0.39±0.01 | -4.6±0.17 | 0.40±0.02 | -4.70±0.17 | 0.33±0.01 | -4.74±0.19 | 0.40±0.01 | -4.67±0.12 |
| **Veins** | 0.41±0.01 | -4.80±0.21 | 0.42±0.02 | -4.79±0.14 | 0.31±0.01 | -4.78±0.19 | 0.43±0.02 | -4.67±0.10 |
| **Aqueduct** | 0.40±0.01 | -4.66±0.25 | 0.40±0.02 | -4.80±0.19 | 0.34±0.02 | -4.51±0.14 | 0.41±0.01 | -4.61±0.22 |

The results of the cross-spectral coherence analyses are summarized in Figure 9. PPG and HRV are seen to exhibit coherence with CSF regions (as well as other brain regions) exclusively at frequencies below 0.2 Hz, while SDPPG exhibit coherence with the rs-fMRI signal at much higher frequencies (closer to 1 Hz). PIR, the metric that is known to reflect low-frequency variations in blood pressure, exhibits coherence with rs-fMRI signals at a much lower frequency (< 0.1 Hz). To quantify these plots, the peak-coherence ridges are plotted for each subject in Figure 10.

**Figure 9.**
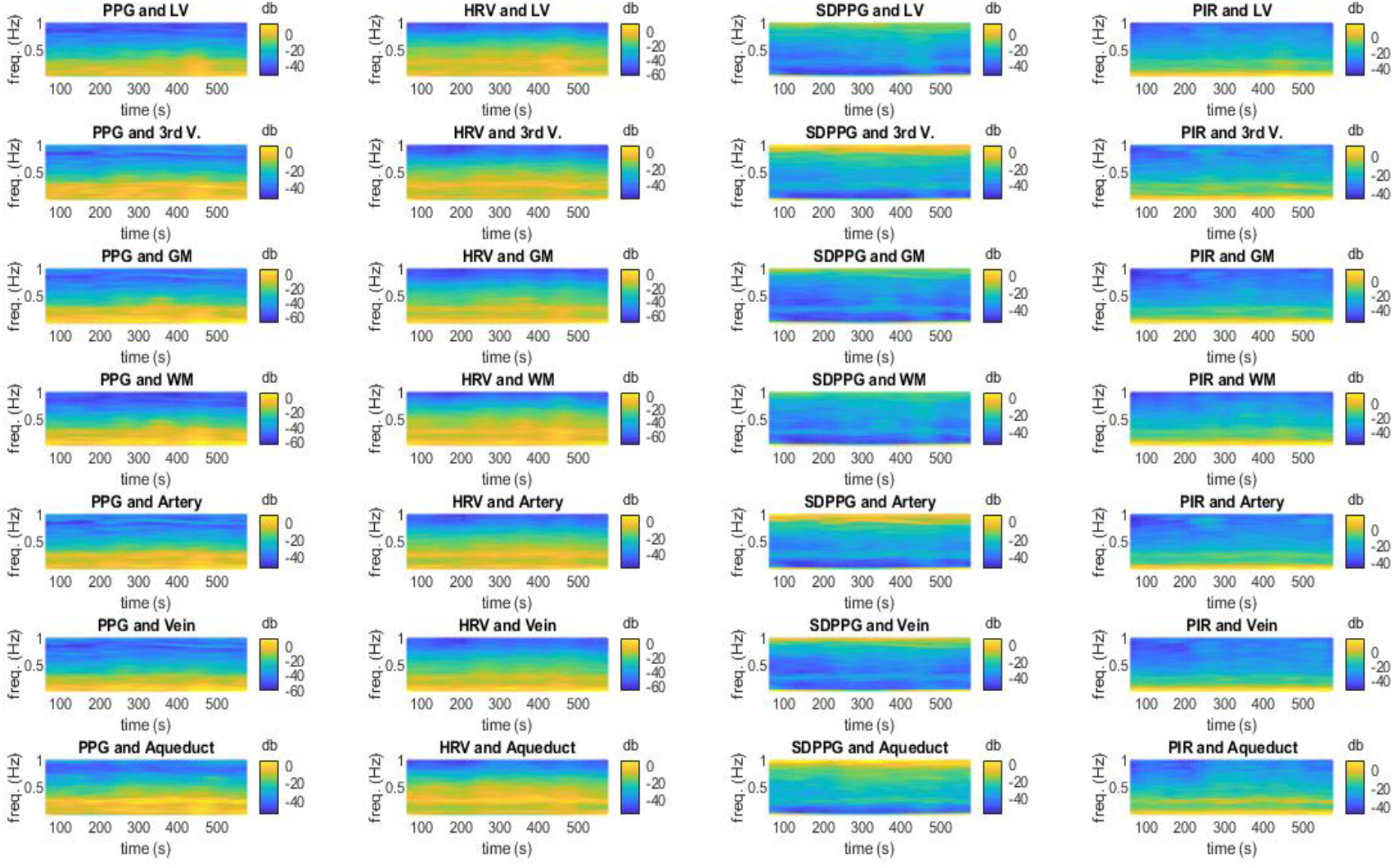
Cross-spectrograms linking PPG-derived signals and fMRI signals in various ROIs. The frequency range is 0.008 to 1 Hz. Coherence is displayed in power per unit frequency (dB/Hz). The cross-spectrograms are generated using the short-time Fourier transform with sliding windows dof 250 s and 90% overlap between steps. High values indicate a high extent shared frequency coupled with high power in these frequencies. The cross-spectrograms have been averaged across all subjects.

**Figure 10.**
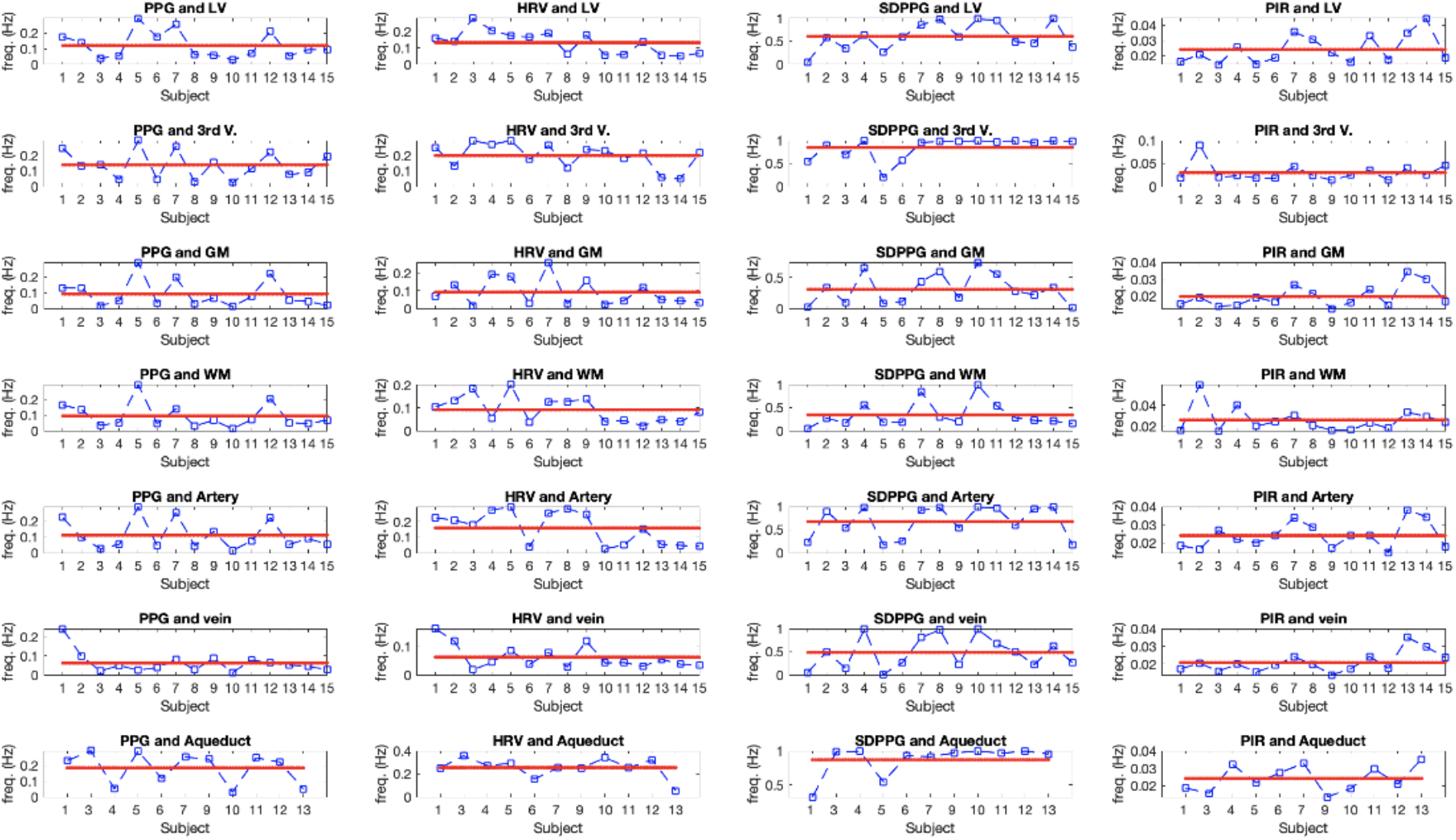
Peak coherent frequencies associating PPG-derived and rs-fMRI signals in various ROIs. Each data point represents one subject. The red line represents the inter-subject average frequency. Frequency values are computed as the weighted average of ridge frequencies in the cross-spectrograms.

The findings in Figure 10 are summarized in Table 3. As shown in Table 3, each PPG-derived parameter exhibits a distinct coherent frequency with rs-fMRI data. Across all frequencies up to 1 Hz, PPG exhibits the strongest coherence with CSF signals near 0.1 Hz, in the range of the M wave and vasomotion. This is similar to the cases of the vascular and tissue ROIs. Conversely, PIR exhibits the strongest coherence with CSF signals in the B-wave range (0.02-0.03 Hz), but this is also found to be the case in the vascular and tissue ROIs. HRV is most strongly coherent with the lateral ventricle at 0.14 Hz, but most coherent with the third ventricle and the aqueduct at higher frequencies (up to 0.2 Hz), similar to the case of the arterial ROI. Interestingly, the HRV coherence is at a much lower frequency in the tissue and venous ROIs. Lastly, for SDPPG, the strongest coherence with all rs-fMRI signals are at a much higher frequency for the CSF regions as well as arterial ROIs (0.6-0.8 Hz), and at a lower frequency for tissue and venous ROIs (0.3-0.4 Hz).

**Table 3.** Summary of peak coherent frequencies between PPG-derived signals and fMRI signals. The frequency range of interest is 0.008-1 Hz. These frequencies are derived from the weighted average of the ridge frequencies extracted from the cross-spectrograms, as shown in Figure 10.

| All values in Hz | PPG | HRV | SDPPG | PIR |
| --- | --- | --- | --- | --- |
| Lateral ventricles | 0.12±0.02 | 0.14±0.02 | 0.61±0.08 | 0.02±0.002 |
| 3rd ventricle | 0.14±0.02 | 0.208±0.02 | 0.84±0.06 | 0.03±0.005 |
| Grey matter | 0.09±0.02 | 0.09±0.02 | 0.31±0.06 | 0.02±0.002 |
| White matter | 0.09±0.02 | 0.09±0.01 | 0.34±0.07 | 0.03±0.003 |
| Arteries | 0.11±0.02 | 0.16±0.03 | 0.68±0.09 | 0.02±0.002 |
| Veins | 0.06±0.01 | 0.06±0.01 | 0.48±0.09 | 0.02±0.002 |
| Aqueduct | 0.19±0.03 | 0.25±0.03 | 0.87±0.07 | 0.02±0.002 |

As shown in Table 4, in the B-wave range, PPG-derived vascular signals account for between 4% and 18% of the rs-fMRI signal. In this frequency range, the PPG signal accounts for the largest CSF signal variance, at ~16%, followed by HRV, which accounts for 13%, and PIR, which accounts for between 11% and 14%. The SDPPG accounts for the least signal variance in CSF in this range, as is also the case in other ROIs.

**Table 4.** Summary of signal variance in rs-fMRI explained by PPG-derived signals. The frequency range of interest is 0.008-0.03 Hz. These frequencies are derived from the weighted average of the ridge frequencies extracted from the cross-spectrograms. In general, each PPG-derived parameter exhibits a distinct coherent frequency with rs-fMRI data.

| Coefficient of determination (r<sup>2</sup>) | PPG | HRV | SDPPG | PIR |
| --- | --- | --- | --- | --- |
| Lateral ventricles | 0.13±0.03 | 0.14±0.02 | 0.07±0.01 | 0.11/0.02 |
| 3rd ventricle | 0.15±0.02 | 0.14±0.03 | 0.11±0.01 | 0.14/0.02 |
| Grey matter | 0.16±0.03 | 0.12±0.02 | 0.05±0.01 | 0.18/0.02 |
| White matter | 0.16±0.03 | 0.11±0.02 | 0.04±0.01 | 0.16/0.03 |
| Arteries | 0.15±0.03 | 0.13±0.03 | 0.07±0.02 | 0.14/0.02 |
| Veins | 0.16±0.03 | 0.13±0.02 | 0.05±0.01 | 0.18/0.03 |
| Aqueduct | 0.10±0.02 | 0.14±0.03 | 0.11±0.02 | 0.11/0.02 |

As shown in Table 5, in the M-wave range, PPG-derived vascular signals account for a similar percentage of the rs-fMRI signal as seen in the B-wave range. In this frequency range, the PPG signal, HRV and PIR all explain similar fractions of rs-fMRI variance in the CSF. The SDPPG accounts for the least signal variance in CSF in this range. Again, the trends in the CSF ROIs are mirrored in the vascular and tissue ROIs.

**Table 5.**
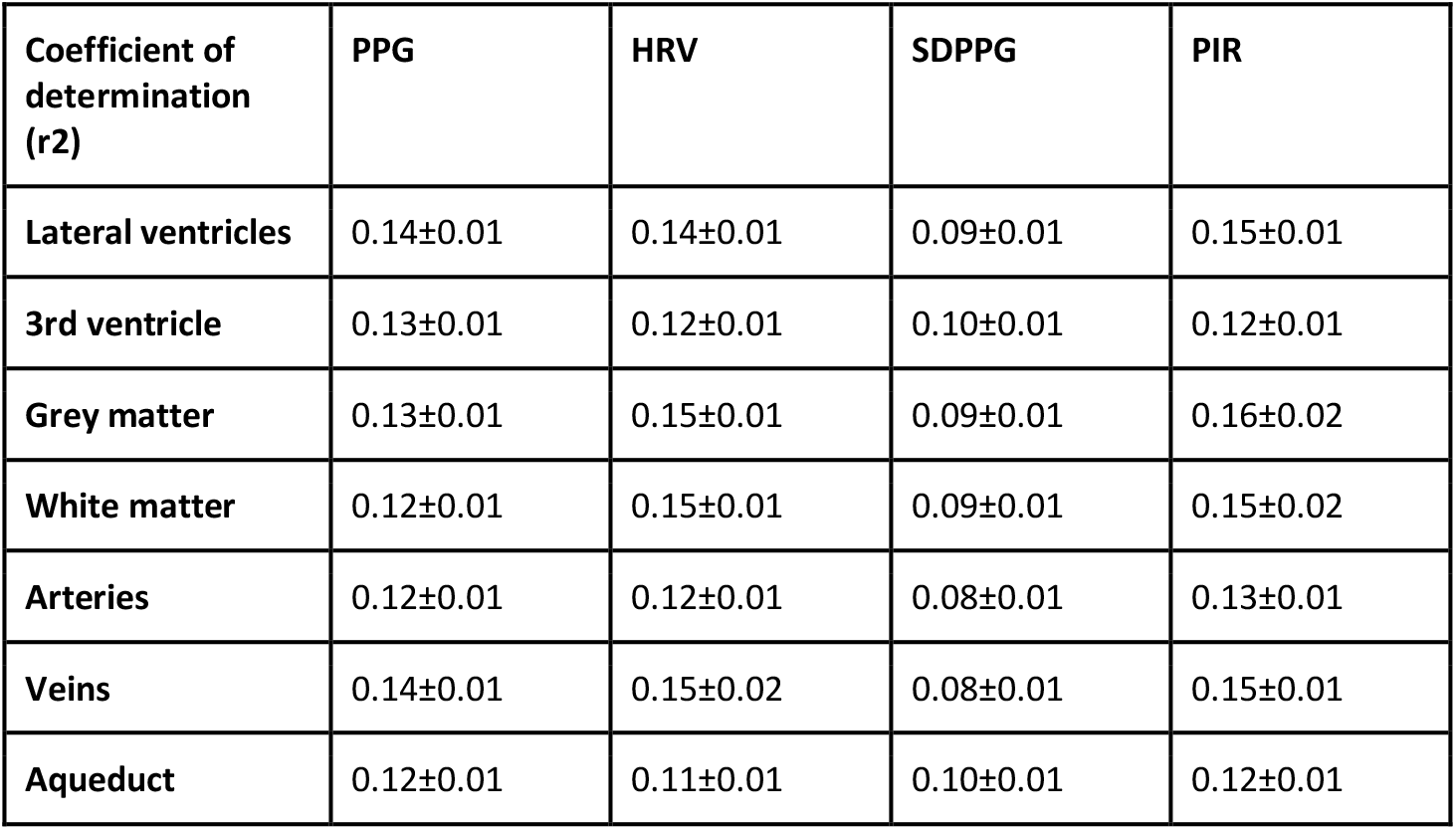
Summary of signal variance in rs-fMRI explained by PPG-derived signals. The frequency range of interest is 0.05-0.15 Hz. These frequencies are derived from the weighted average of the ridge frequencies extracted from the cross-spectrograms. In general, each PPG-derived parameter exhibits a distinct coherent frequency with rs-fMRI data.

## Discussion

Hemodynamic oscillations have conventionally been measured using transcranial Doppler ultrasound (TCD) on a single vessel, most often the internal carotid or the middle-cerebral artery (Schytz et al., 2010). The TCD literature proves that blood pressure related spontaneous CBF fluctuations provide an effective means of characterizing cerebral autoregulation. Rhythmical oscillations in laser Doppler flow have been observed, with characteristic frequencies for the forehead (0.13+/− 0.03 Hz) and finger (0.07+/− 0.02 Hz) (Podgoreanu et al., 2002). Peripheral circulation measured using the photoplethysmograph (PPG) represents a simpler and more common approach to measuring vascular oscillations. It is generally recognized that the high-frequency PPG signal (0.2 - 0.5 Hz) is attributed to respiratory effects, while the lower-frequency component (0.05 - 0.15 Hz) is associated with sympathetic regulation of peripheral vascular resistance (Anschütz and Schubert, 2005; Krupatkin, 2009) as well as the B and M waves (Allan et al., 2018; Bernardi et al., 1996; Middleton et al., 2011). Indeed, PPG has been used to identify a 0.1 Hz power-spectral peak (9 - 27 cycles per minute), which could stem from either vasomotion or the M wave (Kanders et al., 2013; Kiselev et al., 2020). Moreover, the use of ultra-low frequency PPG spectrum for estimating intracranial-pressure (ICP) variations (hence B wave) has recently been demonstrated (Evensen and Eide, 2020; Wagshul et al., 2011). Furthermore, PPG-derived HRV (Claudia Strik et al., 2002) has been extensively studied in terms of its relationship with rs-fMRI functional connectivity (Chang et al., 2009; Shmueli et al., 2007).

It has been suggested that vasomotion is entrained by coordinated oscillations in endothelial calcium concentration (Aalkjær et al., 2011; Mateo et al., 2017), and can propagate in localized regions instead of being systemic (like M waves) (Rayshubskiy et al., 2014). This distinction of course does not rule out possible association between the two physiological phenomena. It has been shown that the amplitude of vasomotion is modulated by arterial blood pressure (Meyer et al., 1988). HRV is constructed from natural variations in the R-R peak intervals, and has been used as a measure of cardiovascular health as well as stress level (Mather and Thayer, 2018; Tsvetanov et al., 2015). Notably, all of these oscillation frequencies are within the range of signal frequencies typically used for resting-state functional connectivity mapping, prompting efforts to clarify their contributions and distinguish them from neurally driven oscillations also situated at <0.1 Hz.

Using ultra-fast MREG fMRI, previous work by Kiviniemi et al (Kiviniemi et al., 2016) demonstrated the vascular contribution to the CSF signal acquired using ultra-fast magnetic resonance encephalography (MREG) sampling at 10 Hz. The Kiviniemi study demonstrated the participation of three frequency bands, namely cardiac (~ 1 Hz), respiratory (~ 0.3 Hz), low-and-very-low frequency (LF 0.023–0.73 Hz and VLF 0.001–0.023 Hz). As a novelty of this study, we are primarily interested in the contributions of low-frequency vascular oscillations (< 0.15 Hz) to the rs-fMRI signal. To further focus our investigation, we interrogate two well-defined vascular oscillatory mechanisms within the low-frequency range, namely the B wave and the M-wave (which overlaps with the frequency of vasomotion). They are both understudied in the MRI literature, but both have established clinical utility (Julien, 2006; Martinez-Tejada et al., 2019). Such low-frequency oscillations are typically challenging to characterize as it is currently impossible to isolate vascular from neuronally driven oscillations in this frequency range. We circumvent these challenges by focusing on CSF regions, which are in theory devoid of neuronal activity.

Moreover, the typical sampling rate (1/TR) of rs-fMRI is too low to allow separation of low-frequency from aliased high-frequency physiological noise. As another novelty of this study, instead of using MREG, we use the more widely available simultaneous multi-slice acceleration EPI to achieve a whole-brain sampling rate of 2.6 Hz, thereby alleviating aliasing from heart beats and respiration. CSF flow fluctuates and reverses direction with cardiac pulsation. The typical CSF flow velocity is 50 mm/s (Zhu et al., 2006). Thus, given the short TR that we use, we expect CSF signals in rs-fMRI to be driven by flow effects. As a further novelty, we use pulse plethysmography (PPG) recordings as an independent reference point for helping to clarify the vascular implications of the CSF signal.

The main findings of this study are: (1) signals in different CSF ROIs are not equivalent in their vascular contributions or in their associations with vascular and tissue r-sfMIR signals; (2) the PPG signal contains the highest signal contribution from the M-wave range, while the PIR contains the highest signal contribution from the B-wave range; (3) in the low-frequency range, PIR is more strongly associated with CSF rs-fMRI signal than PPG itself, and than HRV and SDPPG; (4) PPG-related vascular oscillations only contribute to < 20% of the CSF signal in rs-fMRI, insufficient support for the assumption that low-frequency CSF signal fluctuations directly reflect vascular oscillations.

### Characteristics of different CSF compartments

CSF flows through the lateral ventricles (LV) download into the third ventricle, and thereafter into the cerebral aqueduct, eventually existing into the central canal. Our results show that the third ventricle and the cerebral aqueduct contain more signal at the cardiac frequency than the lateral ventricles (LV) (Figure 2 & 3). In fact, different compartments of CSF exhibit different spectral features altogether. The signal in the lateral ventricle shares greater similarity with low-frequency vascular oscillations, signals in the third ventricle and the aqueduct contain more cardiac pulsatility. Moreover, as shown in Figures 5 and 6, the lateral ventricles contain more low-frequency oscillations of possible vascular origin than the third ventricle and the cerebral aqueduct. Such a distinction is not commonly known in the rs-fMRI community, and can be particularly useful when using the signal averaged over all CSF regions as a regressor for physiological denoising.

As for the difference between the LV, third ventricle and aqueduct, one possibility is the differences in ROI size. The LV is largest and therefore is likely to exhibit the highest SNR. Nonetheless, through-plane flow in the LV is likely more damped compared to that of the other two ROIs, resulting in reduced cardiac pulsatility in the LV signal.

### Associations between CSF signal and vascular and tissue signal

In terms of beat-to-beat dynamics, CSF flow in the aqueduct are well coupled to that of the common carotid arteries (Schmid Daners et al., 2012). On the other hand, alterations in venous dynamics (e.g. through venous compression) are known to alter ICP and lead to reductions in CSF flow (Ichikawa et al., 2018).

The relationships between signals in CSF and non-CSF ROIs in the whole frequency range (0.005-1 Hz) and in the low-frequency range (0.008-0.15Hz) are very similar (Figure 4). Specifically, as we may conjecture from Figures 2 and 3, the CSF signal in the LV is strongly correlated with that in the third ventricle, the arterial ROI, GM and WM. However, the LV signal is only significantly associated with the aqueduct signal in the low-frequency range. Moreover, GM and WM signals are strongly correlated. These are in turn strongly correlated with arterial and venous signals, more strongly in the low-frequency range than in the whole range. In the low-frequency range, signals in the aqueduct are weakly associated with those in the GM, WM and veins, but significantly associated with LV and arterial signals.

The B-wave frequency range contributes < 20% of the fMRI signal from CSF regions (LV, 3rd V and aqueduct). Specifically, the LV and the 3rd ventricle contain 18% and 10% contribution from this frequency range. These values are all in excellent agreement with those measured by Strik et al. (Claudia Strik et al., 2002). The B-wave frequency contributes 40% to the rs-fMRI signal in the large veins, also in agreement with the measurement (of 36%) by Strik et al. However, the contribution of 47% to the grey matter is much higher than reported by Strik et al. for brain parenchyma, possibly due to our segregation of GM and WM (35%). Finally, the B-wave range contributes 23% to the arterial rs-fMRI signal, in agreement with findings by Strik et al. again. The data in Figure 5 hint at greater similarity between arteries and CSF than between veins (and brain tissue) and CSF. The cerebral aqueduct exhibited the lowest fractional signal in the M-wave range, potentially due to the low SNR resulting from the small size of the ROI.

The M-wave frequency band contributes ~28% of the CSF signal in the LV, but only 15% of the signal in the 3rd ventricle. The LV measurement is in excellent agreement with the 27% measured by Strik et al., though in the aqueduct rather than in the LV. In veins, ~25% of the signal fluctuation comes from the M-wave band, lower than the 43% reported by Strik et al. In GM and WM, this figure is 20% and 28%, respectively, and when averaged across the two, is comparable to the 30% measured by Strik et al.. In arteries, a similarly low signal percentage (13%) as in the third ventricle is observed. These data suggest that the arteries share more signal characteristics with the third ventricle and aqueduct rather than with the LV, as reflected in Figure 3 as well.

### Potential implications of PPG-derived signals

It is clear, as seen in Figure 3, that the PPG signal has a clear high-frequency peak (at the cardiac frequency) and a strong low-frequency peak (< 0.5 Hz). The PPG signal has been used as a non-invasive way to directly assess vasomotion (Kanders et al., 2013). The B-wave and M-wave frequency bands contribute ~5% and 26% to the PPG signal (Figure 5 & 6). These figures are consistent with the fact that the B-wave is typically more difficult to observe than the M-wave. Nonetheless, in our study, the PPG provides the most direct method of tracking high- and low-frequency vascular oscillations.

While HRV is often assumed to be an ultra-low-frequency phenomenon in rs-fMRI, only a negligible fraction of the HRV signal is found in the 0.008-0.03 Hz range (Figure 5b). As seen in Figure 3, the power of HRV is mainly distributed between 0.2 and 0.6 Hz, situated between the frequencies of PIR and SDPPG. SDPPG, which is included in this study as a commonly cited measure of arterial compliance, is mainly driven by high-frequency vascular oscillations, presumably reflecting beat-to-beat compliance. PIR, as a surrogate measure of SBP, contains the highest percentage of power in the band < 0.1 Hz, attesting to the relationship between intracranial and systemic blood pressure (Martinez-Tejada et al., 2019).

### Association between CSF signals and PPG-derived signals

PPG measures changes in subcutaneous blood volume that is induced by the pulse pressure wave. This is done by tracking changes in subcutaneous absorption of near-infrared light throughout the cardiac cycle. The PPG signal is measured at the fingertip, which is supplied from the heart via the radial artery to the digital arteries, with a pulse-wave transit time of no more than 0.5 s (Huttunen et al., 2019). The same heart supplies the brain through the aorta and the common carotid artery, with an associated transit time of no more than 0.7 s (Huttunen et al., 2019).

Of all the PPG-derived features, the spectral characteristics of PIR are most similar to that of the LV and the blood vessels, echoing observations in the time domain (Figure 2). On the other end of the spectrum, SDPPG, like the aqueduct, appears to be mainly driven by high frequencies (heart rate), which is also strongly manifested in the aqueduct fMRI signal.

We used short-term cross-correlations via sliding windows as the signals involved are nonstationary. The goal of the cross-correlations is to provide an easily understandable view of the data. In the B-wave range, the PPG signal lags the CSF signal by ~20 s. In a previous report whereby B waves were found to lead arterial blood pressure oscillations by 10 s (Droste and Krauss, 1999), the difference of 10 s in lag can be partially accounted for by the delay between the site of blood-pressure measurement and the fingertip, assuming an immediate relationship between arterial blood pressure and blood flow. A similar delay is found for HRV and SDPPG. However, the peak lag implies that B waves lag PIR, a surrogate for arterial blood pressure. The implications of this finding are unclear.

For the M-wave frequency range, the PPG signal is found to lead the CSF signal by 4-5 s. If indeed Mayer waves are dominating these correlations, then these would likely be driven by the baroreflex (Ghali and Ghali, 2020; Julien, 2006), the time delay of which has been estimated at 2.5 s (Borst et al., 1983), not too distant from our estimate. This time there is no noticeable difference among the various PPG-based metrics in terms of lag and correlation values, and CSF ROIs are similar to non-CSF ROIs in terms of the correlation coefficients. SDPPG accounts for the lowest cross-correlation values, with no distinction between CSF regions and non-CSF regions.

For both the B-wave and M-wave ranges, arteries and the white matter share similar lags, while veins and the grey matter share similar lags with respect to PPG-based signals. Amongst all ROIs, CSF regions also have generally the lowest cross-correlation coefficients with PPG-signals, although the difference is not statistically significant.

While 18% of the LV signal is in the B-wave range (Figure 5), only 13% of the signal variance in this frequency band is explained by variance in the PPG signal (Table 4). Likewise, while 28% of the LV signal is in the M-wave range, only 14% of the signal variance in the M-wave band is explained by variance in the PPG signal. This indicates that a substantial portion of CSF signal fluctuations in rs-fMRI may not directly reflect vascular fluctuations. Indeed, as stated earlier, one of the other signal sources in the B-wave range may be respiratory variability. However, in the M-wave range, it is immediately obvious why 50% of the CSF signal is unaccounted for by PPG. Competing effects from HRV, SDPPG and PIR may account for the difference.

Figure 9 and Table 3 show a clear distinction between the manifestations of the four PPG-based metrics in the CSF signal. Over a broad range up to 1 Hz, PPG exhibits the strongest coherence with CSF signals near 0.1 Hz, in the range of the M wave and vasomotion. Conversely, PIR exhibits the strongest coherence with CSF signals in the B-wave range (0.02-0.03 Hz). This latter frequency range is in contrast with results form a recent study by Whittaker et al., which found arterial blood pressure to the highest degree of fMRI correlations in the 0.06-0.13 Hz range (Whittaker et al., 2019). One likely explanation is that PIR reflects the low-frequency aspect of arterial blood pressure (Ding et al., 2017). Thus, the CSF signal in the 0.02-0.03 Hz range in the LV and third-ventricle could potentially be used as a surrogate for PIR, hence a surrogate for blood-pressure oscillations.

HRV is coherent with CSF signals at close to 0.2Hz, while SDPPG is coherent with CSF signals at 0.6-0.8 Hz. Neither of these are within the range of the oscillations that we focus on in this work (B wave and M wave). In the CSF, coherence frequencies verify slightly between the LV, third ventricle and aqueduct, consistent with our findings of spectral differences across these three compartments. Moreover, the pattern of an artery-CSF coupling, in contrast to a vein-GM coupling, continues. While the coherence frequencies of PPG-fMRI and HRV-fMRI coherences do not differ between CSF and non-CSF regions, in the HRV and SDPPG, the coherence frequencies of arterial and CSF ROIs are similarly high, while the coherence frequencies of veins and GM are similarly low. This observation is consistent with the knowledge that the WM BOLD signal contains more arterial pulsatility effects (Macintosh et al., 2015) while the GM BOLD signal contains substantial large-vein contributions.

### Limitations and future work

Due to the desire to achieve imaging speed, we were limited in spatial coverage and did not include the fourth ventricle. Moreover, non-linear interactions between PPG-related oscillations and CSF signal fluctuations were not explored. These limitations will shape our future work to further explore the vascular origins of the rs-fMRI signal.

